# Defining the genetic architecture of stripe rust resistance in the barley accession HOR 1428

**DOI:** 10.1101/093773

**Authors:** Shaun Clare, William Kitcher, Matthew Gardiner, Phon Green, Amelia Hubbard, Matthew J. Moscou

## Abstract

*Puccinia striiformis* f. sp. *hordei*, the causal agent of barley stripe rust, is a destructive fungal pathogen that significantly affects barley cultivation. A major constraint in breeding resistant cultivars is the lack of mapping information of resistance (*R*) genes and their introgression into adapted germplasm. A considerable number of *R* genes have been described in barley to *P. striiformis* f. sp. *hordei*, but only a few loci have been mapped. Previously, Chen and Line (1999) reported two recessive seedling resistance loci in the Ethiopian landrace HOR 1428. In this study, we map two loci that confer resistance to *P. striiformis* f. sp. *hordei* in HOR 1428, which are located on chromosomes 3H and 5H. Both loci act as additive effect QTLs, each explaining approximately 20% of the phenotypic variation. We backcrossed HOR 1428 to the cv. Manchuria and selected based on markers flanking the *RpsHOR128-5H* locus. Saturation of the *RpsHOR1428-5H* locus with markers in the region found KASP marker K_1_0292 in complete coupling with resistance to *P. striiformis* f. sp. *hordei* and was designated *Rps9*. Isolation of *Rps9* and flanking markers will facilitate the deployment of this genetic resource into existing programs for *P. striiformis* f. sp. *hordei* resistance.

## INTRODUCTION

Barley stripe rust (syn. barley yellow rust) is caused by the obligate biographic fungal pathogen *Puccinia striiformis* f. sp. *hordei. P. striiformis* f. sp. *hordei* is one of the most serious and destructive fungal pathogens affecting barley cultivars worldwide (Line 2002; Yan and Chen 2007a, 2007b; Yan and Chen 2006). *P. striiformis* f. sp. *hordei* was first described in Europe in the late nineteenth century. Periodic epidemics causing substantial yield loss have occurred in multiple regions including the UK and Europe during the 1960s (Dantuma 1964; Macer 1966; Nover and Scholz 1969). A large-scale epidemic in Columbia started with the detection of *P. striiformis* f. sp. *hordei* race 24 in 1975, which is hypothesized to have been introduced from Europe (Dubin and Stubbs 1986; Marshall 1995). The pathogen spread south to Argentina by 1982 and north into Mexico by 1987, Southern USA (Texas) by 1991 and Northern USA (Washington) by 1995 (Chen and Line 1999, 2003; Chen et al. 1995; Line 2002; Marshall 1995; Wellings 2011). Establishment of *P. striiformis* f. sp. *hordei* led to epidemics in South America during 1988 (Sandoval-Islas et al. 1998) and the Pacific Northwest in the USA in 1996 (Line 2002). Today, the pathogen is endemic in Asia, Africa, Europe, and North and South America (Wellings 2011), and is a potentially devastating threat to Australia (Dracatos et al. 2016). The majority of these epidemics have been associated with a sudden transition from the absence or low disease pressure to a rapid proliferation of the pathogen (Dantuma 1964). The impact on barley depends on the environment, the structure of the pathogen population, the use of fungicides, and genetic resistance within cultivars. The durability of genetic resistance is under constant threat by the natural variation and novel mutation in the pathogen and raises the importance of discovering novel sources of resistance.

Genetic resistance is a critical component of integrated disease management and can reduce the economic and environmental impact of chemical fungicides (Chen et al. 1994; Chen and Line 2003; Chen et al. 1995; Dracatos et al. 2016; Yan and Chen 2006). Several screens for genetic resistance have been performed on diverse germplasm including elite, landrace, and wild barley accessions. A total of 26 loci were identified to confer resistance to *P. striiformis* f. sp. *hordei* in barley by Chen and Line (1999) and Nover and Scholz (1969). They found that 21 genes had a recessive mode on inheritance, whereas five had a dominant mode of inheritance (Chełkowski et al. 2003; Chen and Line 1999). To date, five *Rps (Resistance to* Puccinia striiformis) loci have been genetically mapped. *Rps* genes functional against *P. striiformis* f. sp. *hordei* include *rps1* on chromosome 3H (Yan and Chen 2007a), *Rps4* on chromosome 1H (Johnson 1968), and *rps5 (=rpsGZ)* on chromosome 4H (Esvelt Klos et al. 2016; Yan and Chen 2006). *Rps6* on chromosome 7HL and several other colocalizing QTLs condition resistance to several *formae speciales* of *P. striiformis* (Castro et al. 2003; Dawson et al. 2016; Derevnina et al. 2015; Li et al. 2016; Sui et al. 2010; Toojinda et al. 2000).

A major challenge in mapping resistance loci is the presence of multiple *Rps* genes in a genetic background. One such case includes the Ethiopian barley landrace HOR 1428, where Chen and Line (1999) identified two resistance loci: *rpsHOR1428-1* and *rpsHOR1428-2*. To map these genes, we generated an F_2_ mapping population, developed a genetic map, and performed QTL analysis. In parallel, we initiated a backcrossing scheme to isolate individual *Rps* genes for further fine-mapping and positional cloning.

## MATERIALS AND METHODS

### Plant and fungal materials

Seeds for accessions HOR 1428 (PI 548708), Morex (CIho 15773), and Manchuria (CIho 2330) were obtained from the United States Department of Agriculture Germplasm Resources Information Network (USDA-GRIN). HOR 1428 is a 2-row spring Ethiopian landrace, Morex is a 6-row elite spring malting cultivar developed at the University of Minnesota, and Manchuria is a 6-row cultivar spring barley. An F_2_ population was generated from selfing a single HOR 1428 x Morex F_1_ plant. Additionally, a BC_1_ population was generated through backcrossing HOR 1428 x Manchuria F_1_ with the cv. Manchuria as the female. Plants were grown and crossed in the greenhouse with supplemental lighting provided by high-pressure sodium lamps (Philips MASTER SON-T PIA Plus 600W/220 E40). A single individual BC_1_ line was selected to generate a BC_1_F_2_ population. *P. striiformis* f. sp. *hordei* isolate B01/2 was collected in 2001 and were bulked and maintained at the National Institute for Agricultural Botany (NIAB) on susceptible barley cv. Cassata. *P. striiformis* f. sp. *hordei* urediniospores were stored at 6°C after harvesting.

### Pathogen Assays

Four groups of eight seeds were sown in a 1 L pot using a peat-based compost. Plants were grown in a controlled environment chamber at 18°C day and 11°C night using a 16 h light and 8 h dark cycle with lighting provided by metal halide bulbs (Philips MASTER HPI-T Plus 400 W/645 E40) with average light intensity of 5.6 klux. Inoculations were performed on 12-day-old seedlings when the first leaf was fully emerged and prior to the emergence of the second leaf. Inoculum was prepared by suspending fresh urediniospores with talcum powder at a weight ratio of 1:16. Compressed air was used to disseminate inoculum onto seedlings on a spinning platform. After inoculation, seedling pots were sealed in plastic bags and stored in the dark at 8°C to achieve the high humidity required for successful germination of urediniospores. Seedlings were returned to the controlled environment growth chamber after 48-72 h post inoculation.

### Macroscopic phenotyping

Plants were phenotyped macroscopically 12 days post inoculation using the McNeal scale designed for host systems: that ranges from 0 (immune; no visible symptoms) to 9 (completely susceptible; abundant pustule formation, chlorosis absent) (McNeal et al. 1971).

### Microscopic phenotyping

Microscopic phenotyping of *P. striiformis* f. sp. *hordei* leaf hyphal colonization was performed according to Dawson *et al.* (2015). Briefly, after macroscopic phenotyping, leaves were harvested and placed in 15 mL centrifuge tubes containing 1.0 M KOH with a droplet of surfactant (Silwet L-77, Loveland Industries Ltd.). After incubating for 12 h at 37°C, cleared leaves were normalized with three washes of 50 mM Tris at pH 7.5. A final staining solution of 50 mM Tris pH 7.5 contained 20 μg/mL of wheat germ agglutinin conjugated with FITC (WGA-FITC; L4895; Sigma-Aldrich) was incubated overnight, mounted, and visualized under a fluorescent microscope using blue light excitation and a GFP filter. Percent hyphal colonization (pCOL) was estimated by scanning disjoint field of views (FOV) on either side of the longitudinal axis that captured an area of 5.55 mm^2^ (2.72 mm x 2.04 mm). Hyphal spread below 15%, between 15 and 50%, or above 50% of the surface area in the FOV were given scores of 0, 0.5, or 1. pCOL was estimated by summing all scores and dividing by the number of FOV.

### DNA extraction

Approximately 4 cm of second leaf tissue was sampled and placed into 96-well plates. Tissue was lyophilized, pulverized, and then subjected to a CTAB-based DNA extraction protocol adapted for 96-well plate based format. The protocol is a modified protocol from Stewart and Via (1993) that provides PCR-grade genomic DNA (Nick Lauter, personal communication) (Stewart and Via 1993).

### Genetic map construction

SNPs between accessions HOR 1428, Morex, and Manchuria were identified by genotyping with the 1,536 SNPs barley oligonucleotide pool assay (BOPA1) (Close et al. 2009). The HOR1428 x Morex F_1_ was used as a heterozygous control. Genotyping was performed at University of California, Los Angeles Southern California Genotyping Consortium (Los Angeles, CA, USA). Illumina BeadStudio (V2011.1) module Genotyping (v1.9.4) was used to call genotypes based on automatic clustering with manual curation. An iterative approach was used for genetic map construction using the barley consensus genetic map for selecting markers in 10 cM equidistant intervals (Muñoz-Amatriaín et al. 2011). The Sequenom MassPLEX assay was used to genotype 66 markers on the HOR 1428 x Morex F_2_ population and parents. SNPs used in the BOPA1 were extracted in IUPAC format with 100 bp flanking sequence (Supplemental Table 1). Primers were designed using MassARRAY software v3.1, which generates multiplexed pools of up to 32 SNP assays. Sequenom genotyping was carried out at the Iowa State University Genomic Technologies Facility (Ames, IA, USA). Gaps within the genetic map were saturated with 67 KASP (Supplemental Table 2) and 5 CAPS markers (Supplemental Table 3) (Kota et al. 2008). Genotyping using KASP and CAPS markers was performed as described by Dawson *et al.* (2016). JoinMap v4 was used with default parameters and an independence LOD threshold of 4.0 to construct the genetic map of the HOR 1428 x Morex F_2_ population (Supplemental Dataset 1). All genetic distances were estimated using the Kosambi mapping function. Map quality was assessed using recombination fraction plots (Supplemental Fig. 1).

### QTL analysis

Composite interval mapping was performed using QTL Cartographer (v1.17j) model 6 with a 2 cM step size, and a 10 cM window (Basten et al. 1994). A non-redundant genetic map was used for QTL analysis. QTLs were identified using the H_0_:H_3_ model in Eqtl with experiment-wide thresholds calculated with 1,000 permutations with reselection of background markers with a threshold of α < 0.05 (Doerge and Churchill 1996; Lauter et al. 2008).

## RESULTS

### *Puccinia striiformis* f. sp. *hordei* inoculation and inheritance of seedling resistance

Barley accessions HOR 1428, Manchuria, and Morex were challenged with *P. striiformis* f. sp. *hordei* isolate B01/2. Macroscopic phenotyping of first leaves found that HOR 1428 was completely resistant with a McNeal score of 0, whereas Morex and Manchuria were moderately susceptible with McNeal scores of three individual seedlings ranging from 6 to 8 and 5, respectively. To estimate the number of *Rps* genes conferring resistance to *P. striiformis* f. sp. *hordei* isolate B01/2, we inoculated a HOR 1428 x Morex F_2_ population at the first leaf stage. Macroscopic phenotyping of the population found a skewed distribution, with the majority of F2 individuals exhibiting a resistant phenotype with McNeal scores between 0 and 2 (Fig. 1A). Only five individuals showed susceptible phenotypes with McNeal scores between 5 and 8, which reflects the range of symptoms observed on individual Morex seedlings. A single HOR 1428 x Morex F1 seedling included in the screen showed a McNeal score of 2.

**Fig. 1.**
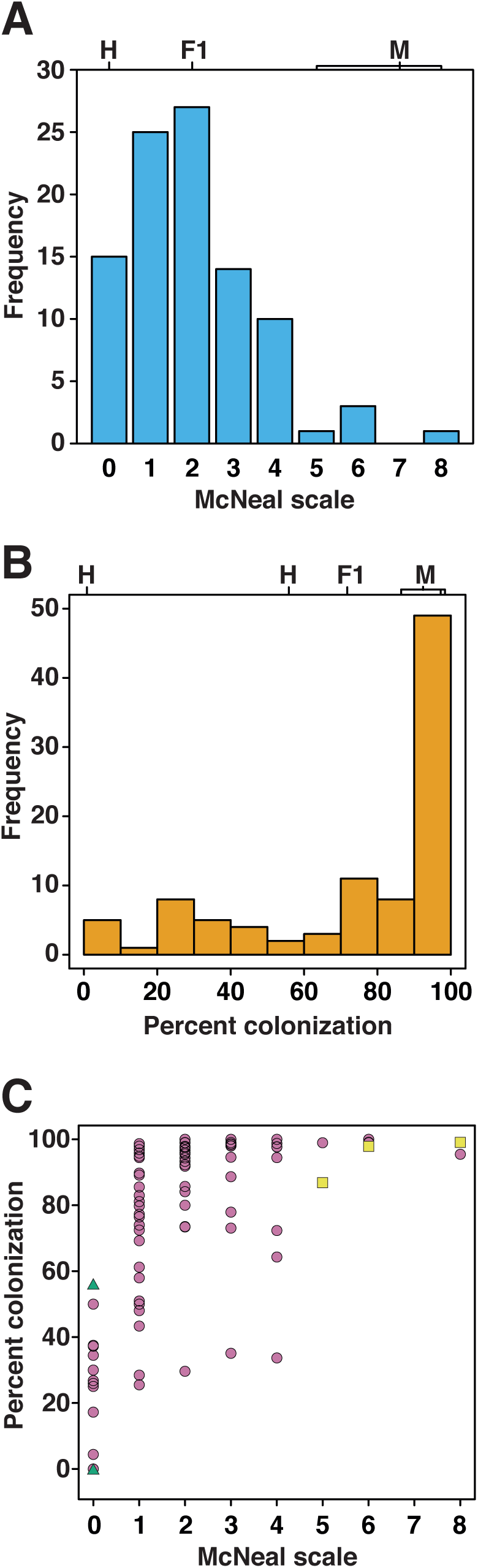
Distribution and two-way plot of disease phenotypes the HOR 1428 x Morex F_2_ population challenged with *Puccinia striiformis* f. sp. *hordei* isolate B01/2. Histograms of (A) McNeal and (B) percent colonization (pCOL) for the F_2_ population with the range of phenotypes for three individual parental (H: HOR 1428 and M: Morex) and a single F_1_ are shown above. (C) Two-way plot of McNeal and pCOL phenotypes, with HOR 1428, Morex, and F_2_ individuals shown as green triangles, yellow squares, and magenta circles, respectively. N = 96 F_2_ individuals.

### Microscopic quantification of *P. striiformis* f. sp. *hordei* hyphal colonization

Previous work established that macroscopic phenotyping does not fully capture the degree of *P. striiformis* hyphal colonization in barley leaves (Dawson et al. 2015). We cleared and stained leaves from HOR 1428, Morex, HOR 1428 x Morex F1, and HOR 1428 x Morex F2 population using WGA-FITC. This stain specifically binds chitin in the fungal cell wall of *P. striiformis* and permits an area-based quantification of hyphal colonization. Average and standard deviations of pCOL for HOR 1428 and Morex were 0.18 ± 0.32 and 0.94 ± 0.07, respectively, whereas the F_1_ had pCOL of 0.73. A highly skewed distribution was observed with approximately half of the F2 individuals, exhibiting near complete (>90%) colonization by fungal hyphae (Fig. 1B). Residual variation was observed in the F2 population that varied from 0 to 90% of leaf area colonized by *P. striiformis* f. sp. *hordei.* A skewed relationship was observed between macroscopic and microscopic phenotypes, with pCOL explaining additional variation in leaves with low McNeal scale (Fig. 1C).

### Quantitative trait locus analysis of barley stripe rust resistance

Parental accessions HOR 1428, Morex, and Manchuria were genotyped using the OPA platform that assays 1,536 SNPs (Close et al. 2009). Based on SNPs that were differential between HOR 1428 and Morex, a panel of Sequenom, KASP, and CAPS markers were developed and assayed on the HOR 1428 x Morex F_2_ population. A genetic map was constructed using 138 markers on nine linkage groups with a total distance of 998.8 cM, with chromosomes 2H and 7H fragmented over two linkage groups (Supplemental Fig. 1). Quantitative trait locus (QTL) analysis was performed with composite interval mapping using macroscopic McNeal and microscopic percent colonization (pCOL) phenotypes. QTL analysis identified two loci mapping to chromosomes 3H and 5H (Fig. 2). The QTLs accounted for 21.5% and 20.9% of the phenotypic variation using the McNeal scale on chromosomes 3H and 5H, with the most strongly linked markers K_2_0566 at 46.74 cM and K_2_0388 at 131.64 cM, respectively (Fig. 2; Table 1). These loci are provisionally designated *RpsHOR1428-3H* and *RpsHOR1428-5H*. Only the 5H QTL was detected using pCOL phenotype and accounted for 41.1% of the phenotypic variation. The peak marker for the QTL was K_1_1090 at 130.32 cM (Fig. 2; Table 1). Both QTLs were contributed by HOR 1428, with the 5H QTL having a slightly stronger additive effect than the 3H QTL, 1.18 versus 0.99 (Fig. 3, Table 1). This observation is coincident with the 5H QTL contributing to a reduction in macroscopic and microscopic resistance, whereas the 3H QTL only contributing significantly to macroscopic resistance (Fig. 2; Table 1).

**Fig. 2.**
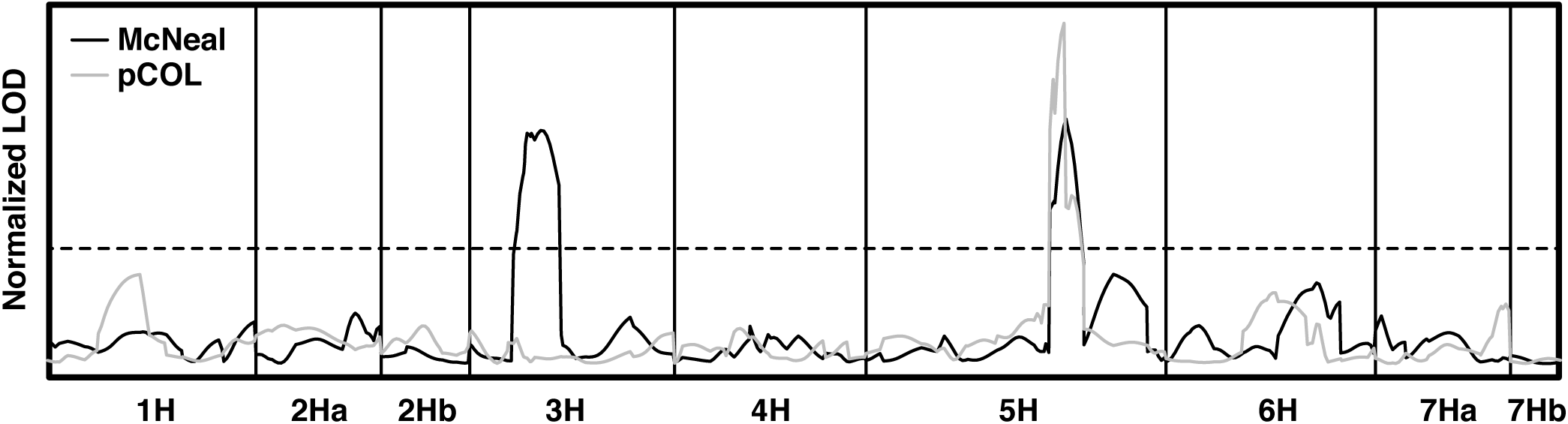
Composite interval mapping of barley stripe rust resistance loci in the HOR 1428 x Morex F_2_ population. LOD scores were normalized to 1.0 based on the significance threshold determined from 1,000 permutation (dashed line). Black and grey solid lines denote the QTL analyses for McNeal and pCOL, respectively.

**Fig. 3.**
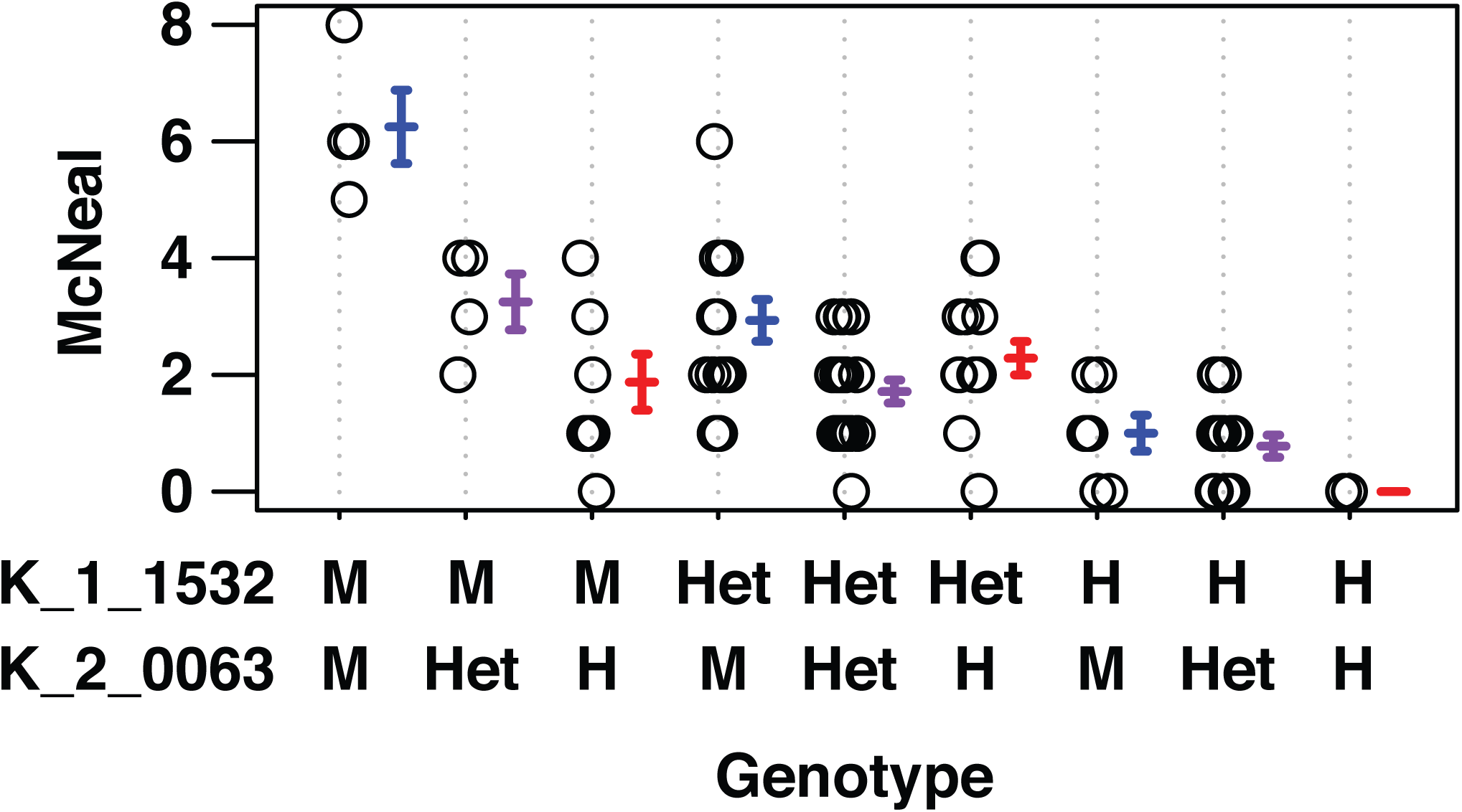
Phenotype by genotype plot of the HOR 1428 x Morex F_2_ population based on peak markers at *RpsHOR1428-3H* and *RpsHOR1428-5H.* Genotypes at markers K_1_1532 and K_2_0063 are denoted as H (homozygous HOR 1428), Het (heterozygous), or M (homozygous Morex).

**Table 1.** Significant resistance QTL identified in the HOR 1428 x Morex F_2_ population challenged with *Puccinia striiformis* f. sp. *hordei* isolate B01/2

| Trait | Chr<sup>a</sup> | cM | Peak Marker | EWT<sup>b</sup> | LOD | AEE<sup>c</sup> | DEE<sup>d</sup> | D/A<sup>e</sup> | PVE<sup>f</sup> |
| --- | --- | --- | --- | --- | --- | --- | --- | --- | --- |
| McNeal | 3H | 46.7 | K_2_0063 | 3.78 | 7.67 | -0.99 | -0.52 | 0.53 | 21.5% |
|  | 5H | 131.7 | K_1_1532 | 3.78 | 8.04 | -1.18 | 0.11 | -0.09 | 20.9% |
| pCOL | 5H | 130.3 | K_1_1090 | 4.04 | 11.96 | -0.25 | 0.16 | -0.63 | 41.1% |

<sup>a</sup> Chromosome
<sup>b</sup> Experiment-wise threshold based on 1,000 permutations with reselection of background markers.
<sup>c</sup> Additive effect estimate
<sup>d</sup> Dominance effect estimate
<sup>e</sup> Ratio of dominance and additive effect estimate
<sup>f</sup> Percent of phenotypic variation explained

### Isolation of *Rps9* in a backcross family

The presence of multiple loci conferring resistance in the HOR 1428 genetic background prompted a backcrossing approach to isolate individual genes from HOR 1428. Manchuria was selected as the recurrent maternal parent as it is susceptible to tested isolates from the United Kingdom for *P. striiformis* f. sp. *hordei* and *P. striiformis* f. sp. *tritici.* KASP markers K_1_1090 at 124.3 cM and K_2_0388 at 143.5 cM were used to select heterozygous BC_1_ individuals flanking *RpsHOR1428-5H,* while counter selecting for homozygosity with a marker at the *RpsHOR1428-3H* locus (K_2_0566 at 37.8 cM). The selected BC_1_ individual was selfed and 94 seeds were grown to fully expanded first leaf and inoculated with *P. striiformis* f. sp. *hordei* isolate B01/2. Saturation of the *RpsHOR1428-5H* locus with nineteen markers in the region found that KASP marker K_1_0292 was in complete coupling with resistance to *P. striiformis* f. sp. *hordei* (Fig. 4). We designate this locus *Rps9*.

**Fig. 4.**
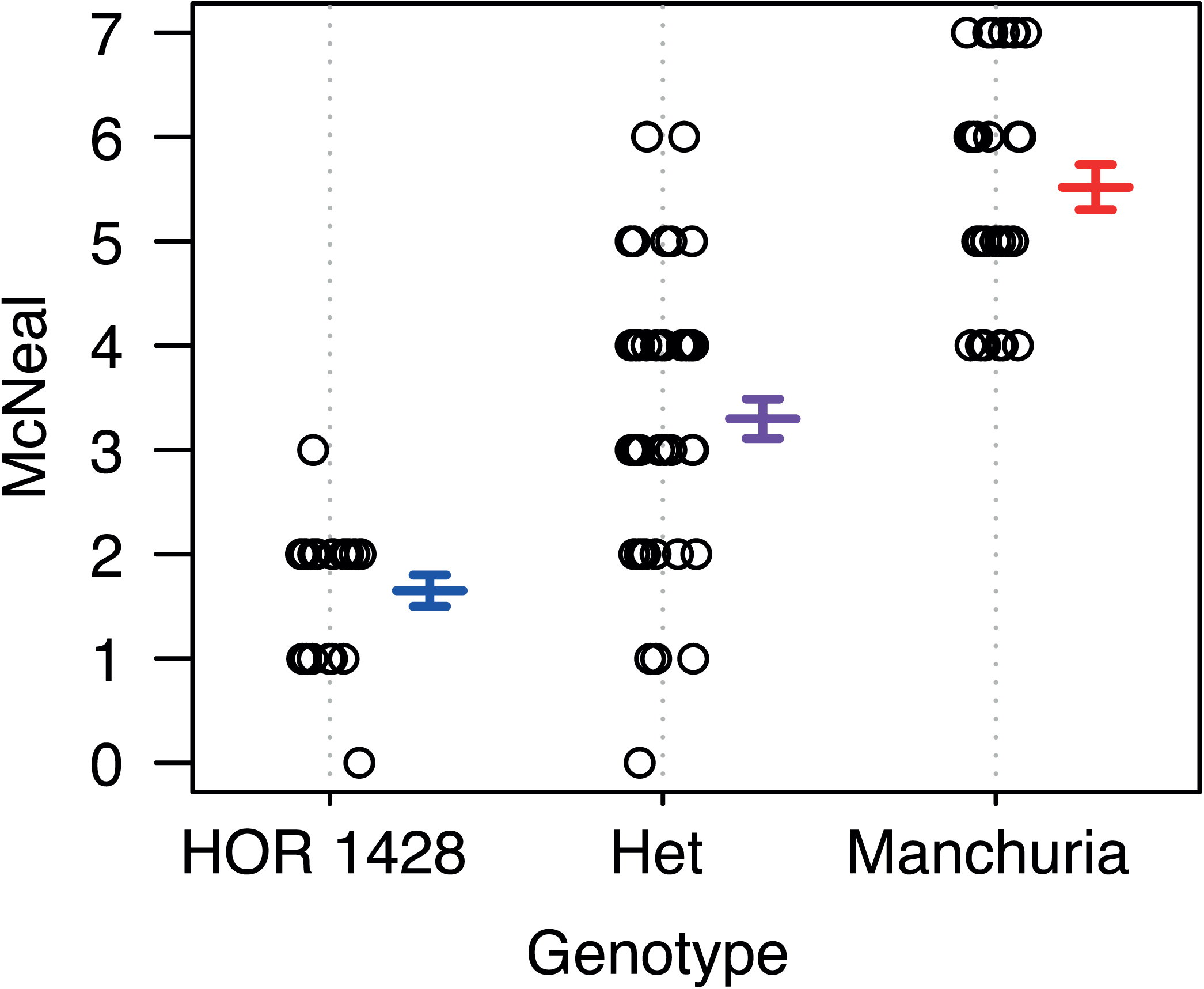
Identification of a cosegregating marker with *Rps9* in a selfed BC_1_ family. Phenotype by genotype plot using 94 BC_1_F_2_ individuals inoculated with *P. striiformis* f. sp. *hordei* isolate B01/2.

## DISCUSSION

Chen and Line (1999) previously identified two recessive loci in accession HOR 1428 conferring resistance to *Puccinia striiformis* f. sp. *hordei*: *rpsHOR1428-1* and *rpsHOR1428-2.* We confirmed the presence of two additive effect QTLs using the HOR 1428 x Morex F_2_ population that mapped to chromosomes 3H and 5H, respectively. We subsequently isolated the *RpsHOR1428-5H* locus present in HOR 1428 in a Manchuria genetic background and designate the locus *Rps9.* The additive nature of *Rps9* suggests that the locus is a semi-dominant resistance gene, which contrasts with the observed recessive nature of *rpsHOR1428-1* and *rpsHOR1428-2* by Chen and Line (1999). The relationship between the loci identified by Chen and Line (1999) and those identified in this study is unclear, but the development of isogenic backcrossed material will facilitate inheritance and specificity of these genes with diverse isolates of *P. striiformis* f. sp. *hordei.*

Discovery of novel sources of resistance is crucial to combat the threat of *P. striiformis* f. sp. *hordei* on barley due to ongoing erosion of resistance genes in cultivation (Castro et al. 2003). Introgression and deployment of resistance loci in elite cultivars requires careful management to minimize any breakdown in resistance. This work sets out to exploit the substantial reservoir of resistance loci present within landrace barley accessions, such as accession HOR 1428, as transferring to adapted germplasm. This is crucial to minimize both outbreaks in endemic locations and preempt potential geographic expansion into regions where *P. striiformis* f. sp. *hordei* has not been recorded, such as Australia (Derevnina et al. 2015; Dracatos et al. 2016). Isolation of additional novel sources of resistance will expedite breeding of gene stacks functional against *P. striiformis* f. sp. *hordei* (Castro et al. 2003; Figueroa et al. 2016; McDonald 2014). The isolation of *Rps9* and markers flanking the interval will facilitate the deployment of this genetic resource into existing programs for barley stripe rust resistance.

HOR 1428 is highly resistant to the nonhost pathogen *P. striiformis* f. sp. *tritici,* the causal agent of wheat stripe rust, a devastating pathogen of wheat (Dawson et al. 2015). This raises the important question of whether HOR 1428 harbors the same genetic architecture to alternative *formae speciales* of the pathogen, and more specifically whether *Rps9* possesses functional overlap against *P. striiformis* f. sp. *tritici?* The isolation of *Rps9* in a susceptible genetic background of Manchuria and future development of a near-isogenic line will permit the testing of this hypothesis, as Manchuria is susceptible to all tested UK isolates of wheat stripe rust (Dawson et al. 2015). In addition, this genetic material will facilitate the future fine mapping, candidate gene identification, and cloning of the gene underlying *Rps9*.

## ACKNOWLEDGEMENTS

We are grateful to the horticulture staff at the John Innes Centre for maintaining excellent plant growth facilities, Richard Goram at the John Innes Centre genotyping facility for KASP genotyping, Inmaculada Hernández-Pinzón for laboratory management, and Rosemary Bayles and Colwyn Thomas for constructive scientific discussions. We appreciate the access to Sequenom genotyping at the Iowa State University Genotyping Facility (Ames, Iowa, USA). Lastly, we greatly appreciate the free, open, and quick access to barley seed from USDA-GRIN (Aberdeen, Idaho, USA). Funding for this work was contributed by Biotechnology and Biological Sciences Research Council (BB/J004553/1) and the Gatsby Charitable Foundation.

## SUPPLEMENTAL MATERIALS

**Supplemental Figure 1.** Recombination fraction plot for HOR 1428 x Morex F_2_ population genetic map.

**Supplemental Table 1.** Sequence used in Sequenom marker development for constructing the HOR 1428 x Morex F_2_ genetic map.

**Supplemental Table 2.** KASP markers used in developing the HOR 1428 x Morex F_2_ genetic map.

**Supplemental Table 3.** CAPS markers used in developing the HOR 1428 x Morex F2 genetic map.

**Supplemental Dataset 1.** Genetic map of HOR 1428 x Morex F_2_ population inoculated with *P. striiformis* f. sp. *hordei.*

